# GRNITE: Gene Regulatory Network Inference with Text Embeddings

**DOI:** 10.1101/2025.11.25.690454

**Authors:** Ali Azizpour, Narein Rao, Santiago Segarra, Luay Nakhleh, Nicolae Sapoval

## Abstract

Gene regulatory networks (GRNs) capture complex regulatory relationships that govern gene expression in cells. Inference of GRNs from single-cell RNA-seq (scRNA-seq) data has been an active topic of research in the past several years. However, despite the improvements in the data quality, the GRN inference problem remains a challenging task with many approaches showing variable performance dependent on the organism and cell type. To improve the quality of GRN inference and enable more comprehensive exploratory analyses of GRNs across various phenotypes of interest, we developed a two-stage meta-method called GRNITE. In the first step, GRNITE leverages LLM-based embeddings of plain text gene descriptions to create a prior gene interaction graph, which is then optimized with a graph neural network (GNN) to achieve a “universal” biological prior for GRN inference. In the second step, GRNITE uses a GNN to incorporate information from a GRN inferred from scRNA-seq data with any baseline inference method into our prior. The result of this two-step approach is a near-universal improvement in AUROC and recall of all evaluated methods, with minor trade-offs in precision. Furthermore, GRNITE is a lightweight meta-method, which adds a minimal amount of extra compute time on top of the original GRN inference performed. GRNITE and our pre-trained universal prior GRN are available on GitHub: https://github.com/aliaaz99/GRNITE.

## 1 Introduction

Gene regulatory networks (GRNs) represent complex molecular interactions that govern gene expression within cells. Inference of GRNs from gene expression data is a longstanding problem in computational biology with multiple methods developed over the past two decades [11,12,22,8,23]. With the emergence of single-cell sequencing technologies, methods for GRN inference from these data began to be developed. However, due to current technological limitations, single-cell RNA sequencing (scRNA-seq) data is prone to gene dropout [38]. This challenge encouraged the development of methods that can explicitly account for the noisy nature of data, as well as the development of multi-omic methods that aim to leverage multiple data modalities (e.g., scRNA-seq + scATAC-seq) to improve the resolution and robustness of GRN inference [31,19,21].

Recently, deep neural networks, such as convolutional neural networks (CNNs) and variational autoencoders (VAEs), have been applied to scRNA-seq data to model complex, non-linear regulatory relationships [32,6,25]. While effective, these methods mostly focus on pairwise transcription factor (TF)–target interactions and often overlook the global regulatory structure of GRNs. Graph neural networks (GNNs) [29] offer a powerful alternative by explicitly modeling genes as nodes and regulatory relationships as edges. GNNs have been successfully used in diverse domains such as social [30,15,24], biological [18,3,17,5,28], and wireless communication networks [34,35] to model structured interactions between entities, enabling tasks such as node classification, community detection, and link prediction [9]. Prior work [27,16] has demonstrated that casting GRN inference as a link prediction problem with a graph autoencoder allows effective integration of global regulatory structure and improves robustness to noise in single-cell data.

Notably, in parallel to these developments, databases of ChIP-seq-confirmed TF-target interactions were expanded, allowing for the construction of stronger evidence-based priors for GRN inference [20,36,2]. These enriched resources have supported the emergence of prior-augmented GRN inference frameworks. Despite these advances, there remains a need for robust and reliable GRN inference approaches that can be applied in exploratory studies with limited references [26]. With the expanded knowledge of individual gene functions, databases describing those in natural language also grew [4]. Additionally, with the emergence of large language models (LLMs) as powerful zero-shot knowledge extractors for natural language, automated knowledge graph construction from plain text became widely accessible [7,37].

Based on a recent benchmarking study of the effectiveness of various pretrained gene embeddings for downstream biological tasks [37], GenePT [7] was identified as a high-quality embedding model for gene-level features. This finding suggests that gene embeddings derived from plain text descriptions can be leveraged as useful priors for gene-based biological inference tasks. Thus, our goal in this study is two-fold: (1) we evaluate a hypothesis that text-based gene embeddings can be used as an effective prior for GRN inference, and (2) we develop GRNITE, a GNN-based tool that can be used in conjunction with any scRNA-seq GRN inference software in order to gain improved performance for GRN inference.

In particular, our framework for **GRN I**nference with **T**ext **E**mbeddings (GRNITE) leverages text modality by replacing co-expression priors with a text-informed prior derived from gene descriptions. This prior is then denoised and refined using curated TF–target interactions to form a biologically grounded scaffold. Moreover, rather than proposing another standalone GRN inference algorithm, we design GRNITE as a flexible pipeline: our enhanced prior is integrated with the output of any existing GRN inference method, allowing us to systematically improve performance and recover biologically plausible interactions without replacing the underlying method itself.

Our results demonstrate that GRNITE increases the Jaccard similarity between the inferred and ground-truth edge sets on benchmarking data. Additionally, we show improvement in balanced accuracy and a universal increase in recall across benchmarking datasets. Furthermore, GRNITE increases inter-method consensus and improves the accuracy of consensus edge sets. Finally, we demonstrate that on biological data, GRNITE enables the superior recovery of key TF-target interactions.

## 2 Methods

In this section, we present **GRNITE**, our method for enhanced GRN inference using text-derived embeddings. An overview of our three-step framework is shown in Figure 1. In Section 2.1, we describe how to construct a text-informed prior graph. Next, in Section 2.2, we apply a graph autoencoder to further refine this prior using additional biological knowledge into a unified prior graph. In Section 2.3, we design a pipeline that leverages this improved prior to enhance the output of any arbitrary GRN inference method. Finally, in Section 2.4, we describe the evaluation setup, including datasets, metrics, and experimental procedures.

**Fig. 1:**
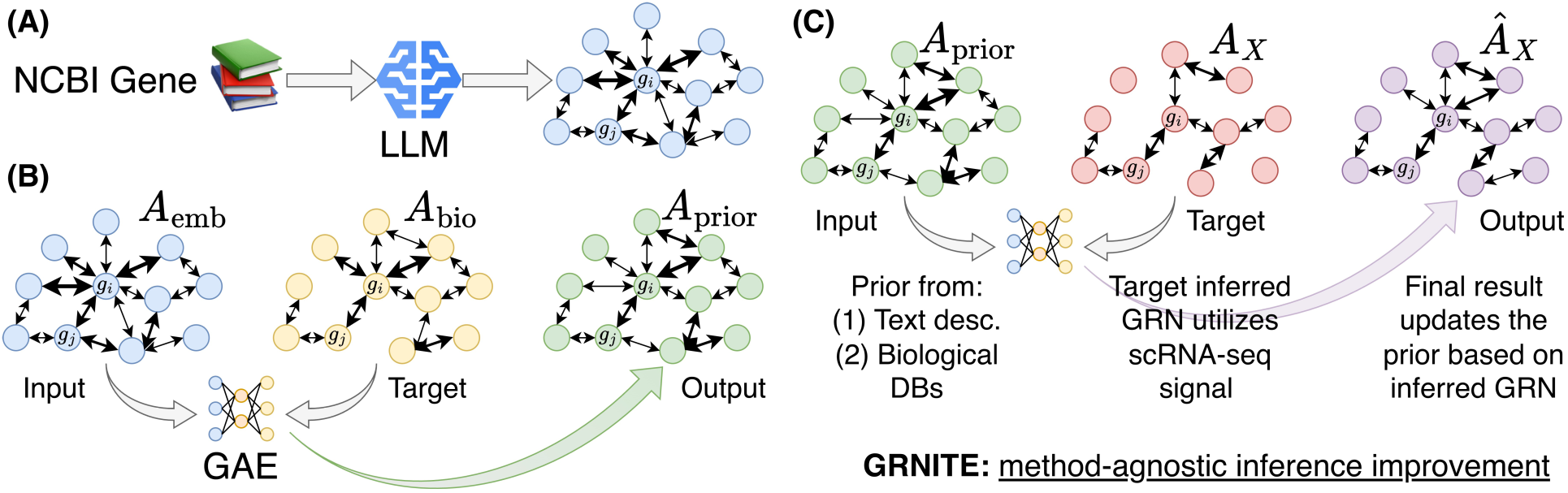
Overview of the GRNITE framework. (A) Embeddings for gene descriptions from NCBI Gene database are computed with a large language model (LLM), and cosine similarity between the resulting vectors is used to construct an embedding prior adjacency matrix (**A**_emb_). (B) A graph autoencoder (GAE) model is trained to reconstruct a biological prior graph (**A**_bio_, based on a database of known TF-target interactions) from **A**_emb_. The final output is an updated prior graph **A**_prior_ that fuses information from text embeddings and known interactions. (C) At inference time, a GRN **A***_X_* is inferred with an arbitrary method from scRNA-seq gene expression data. Then a GAE is trained to reconstruct **A***_X_* from **A**_prior_, and the final prediction **Â** *_X_* is produced.

### 2.1 Text-informed Prior Construction

We begin by constructing a prior gene interaction graph in which connections reflect similarities in text-derived information. To achieve this, we first obtain a text-based embedding vector for each gene by following a procedure similar to the GenePT approach [7]. Specifically, we first obtain gene information and text summaries from the NCBI Gene database. Then we filter out genes that do not have a matching taxonomic ID for our dataset of interest, and any genes that have an empty gene symbol. Finally, the “Summary” field containing brief summary description of the gene, its function (whenever known) and products in plain-text English [4], from the downloaded database is used to compute gene embeddings with Qwen3 embedding 8B model [33]. This results in a **X**_emb_ ∈ R*^G^*^×^*^F^* matrix of gene embeddings where *G* is the number of genes, and *F* = 4096 by default.

Now, let **x***_i_* = **X**_emb_(*i,* :) ∈ R*^F^* denote the embedding of gene *i*. Using these embeddings, we compute a weighted similarity matrix **W**_emb_ ∈ R*^G^*^×^*^G^* by measuring the cosine similarity between pairs of genes:

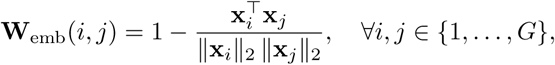

This matrix quantifies how close two genes are in the text-embedding space, with smaller values indicating greater similarity.

Next, we convert this weighted similarity matrix into a binary adjacency matrix **A**_emb_ ∈ {0, 1}*^G^*^×^*^G^*. To do so, we threshold the distances using the average pairwise distance across all gene pairs:

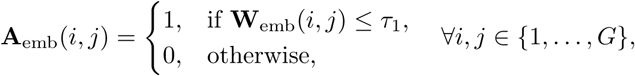

where *τ*_1_ is a tunable hyperparameter. This retains only the most similar gene pairs, forming a sparse prior graph that captures strong relationships based on textual information.

### 2.2 Biology-informed Prior Enhancement

In this step, our goal is to integrate biological knowledge, such as experimentally validated TF–target interactions, into the text-derived graph **A**_emb_ to construct a refined and biologically meaningful prior graph. Let **A**_bio_ ∈ {0, 1}*^G^*^×^*^G^* denote the biological prior adjacency matrix, where **A**_bio_(*i, j*) = 1 if a known interaction between genes *i* and *j* exists in curated databases. Our objective is to learn a new prior graph **A**_prior_ that combines the information in **A**_emb_ and **A**_bio_, i.e., combines dense text-based similarities with high-confidence biological interactions.

To achieve this, we employ a graph autoencoder (GAE) that takes **A**_emb_ as the structural input and uses **X**_exp_ as node features, where **X**_exp_ ∈ R*^G^*^×^*^C^* is the gene expression matrix with *C* denoting the number of cells. Formally, we denote the encoder as a function 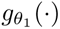 parameterized by *θ*_1_ and the decoder as 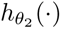 parameterized by *θ*_2_. The encoder maps node features into a latent representation:

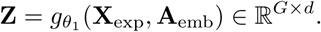

In practice, 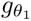 consists of multiple topology-aware graph convolution (TAGConv) layers [10]. A single TAGConv layer is defined as:

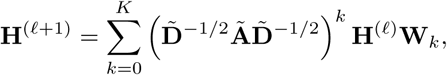

where 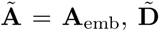 is its degree matrix, **H**^(*ℓ*)^ is the input feature matrix at layer *ℓ*, **W***_k_* are learnable weights, and *K* is the number of hops from which the data is aggregated. Unlike standard GCN layers [14] that aggregate information only from one-hop neighbors (*K* = 1), TAGConv aggregates information from multi-hop neighborhoods up to *K* hops, enabling richer propagation of topological information and better capturing long-range dependencies in the prior graph. The encoder initializes with **H**^(0)^ = **X**_exp_ and produces latent representations **Z** = **H**^(*L*)^ ∈ R*^G^*^×^*^d^* after *L* layers.

Given the latent embeddings **z***_i_* for each gene, the decoder reconstructs an edge score using a bilinear form:

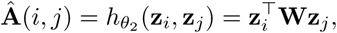

where **W** ∈ R*^d^*^×*d*^ is a learnable weight matrix.

To train the model, we compare the reconstructed scores **Â** against the biological prior adjacency matrix **A**_bio_. However, since **A**_bio_ is highly sparse, directly evaluating the loss over all *G*^2^ possible pairs would lead to severe class imbalance and redundant computations. Therefore, we perform *negative sampling* : we include all positive edges (where **A**_bio_(*i, j*) = 1) and a randomly sample a subset of negative edges (where **A**_bio_(*i, j*) = 0) from the upper triangular part of the matrix. Let *Ω* denote the set of sampled edge pairs.

The reconstruction loss is then defined as:

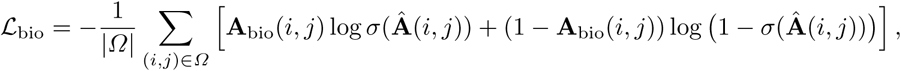

where *σ*(·) denotes the sigmoid activation applied to the raw predicted interaction logits. A positive weighting term can optionally be applied to further mitigate class imbalance by emphasizing true interactions.

The optimal parameters are obtained by minimizing this loss:

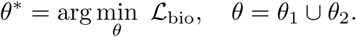

After training, the final reconstructed edge probabilities are obtained as:

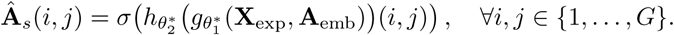

Finally, the refined prior graph is constructed by thresholding the reconstructed scores:

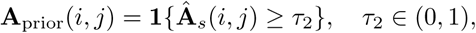

where *τ*_2_ is a tunable threshold. In this way, we first learn the optimal encoder and decoder parameters using **A**_bio_ as supervision, and then use the trained model to produce refined edge scores that are subsequently binarized to construct the final prior graph.

This procedure retains biologically supported edges while preserving additional plausible interactions inferred from text. Since the encoder operates on **A**_emb_, genes connected in the text-derived graph naturally acquire similar embeddings; however, by supervising reconstruction using **A**_bio_, the model is encouraged to suppress edges that are inconsistent with biological evidence, effectively denoising and refining the text-based prior.

### 2.3 Integration with Existing GRN Inference Methods

In the final step, we incorporate the enhanced prior graph obtained from the previous stage to improve the output of any arbitrary GRN inference method. Since this prior embeds complementary information from both biological databases and text-derived embeddings, our goal is to leverage it in a way that enhances, rather than replaces, existing GRN inference pipelines.

Let **A**_X_ ∈ [0, 1]*^G^*^×^*^G^* represent the output of an arbitrary GRN inference method *X*, obtained from the gene expression data **X**_exp_. We seek to refine this output graph by incorporating the rich contextual knowledge captured in the prior graph **A**_prior_.

To achieve this, we employ a graph autoencoder similar to the previous step.

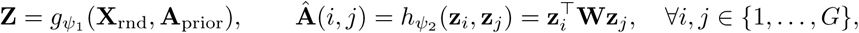

where **Z** ∈ R*^G^*^×^*^d^* is the latent gene representation, **W** ∈ R*^d^*^×^*^d^* is a learnable bilinear matrix. Unlike the previous step, where expression data **X**_exp_ was used as node features, here we use random Gaussian features **X**_rnd_ to prevent information leakage from the target method **A**_X_, which is already inferred from expression data [1]. The encoder is constructed using TAGConv layers, which enable multi-hop message passing and effectively capture long-range dependencies encoded in **A**_prior_.

Different from the previous step, the training objective here encourages the reconstructed graph **Â** to remain consistent with both the target graph **A**_X_ and the enhanced prior **A**_prior_. To accomplish this, we again employ negative sampling: we include all positive edges (where **A**_X_(*i, j*) = 1) and a randomly sampled subset of negative edges (where **A**_X_(*i, j*) = 0) from the upper triangular part of the matrix to avoid duplication. Let *Ω* denote this set of sampled edge pairs.

The two binary cross-entropy with logits losses are defined as:

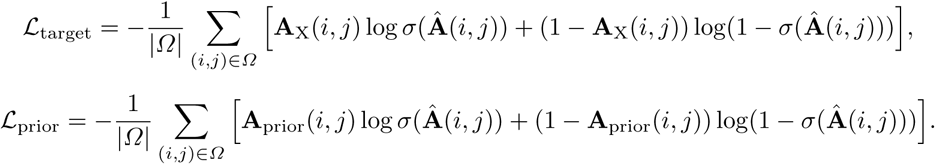

The total loss combines these two objectives as:

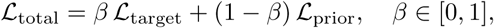

where *β* balances the emphasis between maintaining fidelity to the target graph and preserving similarity to the enhanced prior. The optimal parameters are obtained as:

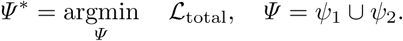

Once trained, the final reconstructed scores are computed using the optimized model:

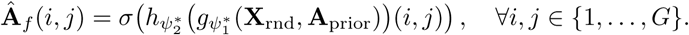

Finally, we threshold these reconstructed scores to obtain the refined GRN:

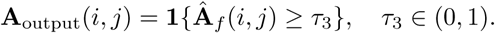

This formulation allows GRNITE to remain fully agnostic to the underlying GRN inference algorithm. The encoder–decoder operates over the prior graph **A**_prior_, ensuring that the latent representations already incorporate biologically grounded and text-informed relationships, while the loss function encourages consistency with the chosen target method. Consequently, this approach enhances the target graph by pruning weak or inconsistent edges and, more importantly, by recovering biologically plausible edges that the base method might have missed. Since the model is independent of the specific choice of X, any GRN inference method can serve as the target, allowing flexible and systematic improvement of existing GRN pipelines.

### 2.4 Datasets and Evaluation

#### Datasets

We leveraged commonly used benchmarking datasets for GRN inference evaluation. In particular, we employed two human and five mouse datasets from BEELINE considering both TFs and 500 and 1000 highly variable genes (HVGs), refered to as BEELINE TF+500 and BEELINE TF+1000 data. We also used simulated data from GRouNdGAN [39]. For BEELINE we relied on accompanying ChIP-seq data to provide ground truth reference for interactions, while for GRouNdGAN we used the ground truth GRN used for the simulation. For the analysis of biological single-cell data, we used two datasets, a high-fat-diet mouse pancreatic *β*-cell dataset (HFD) and peripheral blood mononuclear cell dataset (PBMC). The high-fat-diet mouse pancreatic *β*-cell dataset is divided into two subsets: cells with high *Cd81* expression and cells with low *Cd81* expression. Four immune cell-type specific datasets were also derived from the peripheral blood mononuclear cell dataset, namely classical monocytes, myeloid dendritic cells, naive B cells, and naive CD4 T cells. We acquired 27 ground truth driver TF-target interactions for the high-fat-diet mouse dataset following the workflow provided in KEGNI [16]. We also acquired 1788 TF-target interactions for all immune cell-type specific interactions within the PBMC dataset curated from top 100 TF-target interactions available via cistromeDB [20,36].

#### Evaluation

GRN inference methods are commonly evaluated using metrics such as the Area Under the Receiver Operating Characteristic curve (AUROC) and the Jaccard index. In line with prior work and the BEELINE benchmark [23], we adopt the same evaluation metrics across all datasets to ensure a fair comparison. The evaluation is performed over the entire space of possible TF–target interactions, considering all potential TF–target pairs as candidates for regulatory edges.

Following the BEELINE evaluation setup, the AUROC in our case is equivalent to the balanced accuracy. This is due to the ROC curve being reduced to three operating points: the two trivial cases of predicting all edges as positive or all as negative, and the operating point corresponding to the inferred network. The area under this curve is equal to the average of the *true positive rate* (TPR) and the *true negative rate* (TNR), i.e., AUROC = (TPR + TNR)*/*2, where TPR = TP*/*(TP + FN) measures the proportion of correctly predicted true edges, and TNR = TN*/*(TN + FP) measures the proportion of correctly predicted non-edges. This interpretation is particularly meaningful for GRNs as true regulatory interactions are sparse, resulting in a strong class imbalance.

In addition to AUROC, we report the Jaccard index to directly measure the overlap between the predicted and reference edge sets. For a predicted edge set *Ê* and ground truth edge set *E*, the Jaccard index is defined as: Jaccard(*Ê, E*) = |*Ê* ∩ *E*|*/*|*Ê* ∪ *E*|. While AUROC quantifies how well positive edges are ranked above negative ones, the Jaccard index evaluates exact edge-set similarity, making it a complementary and stricter measure for GRN assessment.

Finally, since in practice no ground truth is available, and consensus edges between multiple method runs are often used as a proxy for high confidence interactions, we also evaluated Jaccard similarities of edge sets inferred by different methods. We further examined how this overlap in edge predictions is affected by application of GRNITE, and whether the updated edge set intersection offers a better AUROC when compared to the baseline.

## 3 Results

### 3.1 Baseline performance

We evaluate the baseline performance of the methods used for inference of the target GRN from the scRNA-seq data on BEELINE and GRouNdGAN datasets (Tab. 1). First, we note that CellOracle has the highest AUROC on BEELINE, and highest Jaccard similarity to true edge sets across all data. On the GRouNdGAN dataset PORTIA achieves the highest AUROC, with CellOracle and GRNBoost being second best in performance. DAZZLE [38] was unable to run on GRouNdGAN simulated data, and larger biological datasets, and hence has been only included in the BEELINE comparisons.

**Table 1:** Baseline performance of the GRN inference methods measured as the mean (± st. dev.) AUROC and edge set Jaccard similarity for the BEELINE and GRouNdGAN datasets. Highest mean value in each column is underlined.

| Method | AUROC |  |  | Jaccard Sim. |  |  |
| --- | --- | --- | --- | --- | --- | --- |
|  | BEELINE (500) | BEELINE (1000) | GRouNdGAN | BEELINE (500) | BEELINE (1000) | GRouNdGAN |
| Correlation | 0.495 ± 0.028 | 0.498 ± 0.028 | 0.516 ± 0.035 | 0.015 ± 0.011 | 0.011 ± 0.008 | 0.019 ± 0.008 |
| GRNBoost | 0.502 ± 0.017 | 0.504 ± 0.011 | 0.591 ± 0.066 | 0.016 ± 0.009 | 0.011 ± 0.007 | 0.042 ± 0.019 |
| PORTIA | 0.501 ± 0.012 | 0.502 ± 0.005 | <u>0.608</u> ± 0.063 | 0.015 ± 0.008 | 0.010 ± 0.005 | 0.068 ± 0.033 |
| DAZZLE | 0.523 ± 0.021 | 0.526 ± 0.025 | — | 0.017 ± 0.009 | 0.012 ± 0.007 | — |
| CellOracle | <u>0.690</u> ± 0.048 | <u>0.697</u> ± 0.050 | 0.593 ± 0.029 | <u>0.112</u> ± 0.059 | <u>0.115</u> ± 0.063 | <u>0.125</u> ± 0.020 |
| SCENIC | 0.510 ± 0.005 | 0.506 ± 0.004 | 0.514 ± 0.016 | 0.019 ± 0.010 | 0.012 ± 0.008 | 0.027 ± 0.031 |

### 3.2 Improvement in inference performance

#### BEELINE Datasets

We evaluated the improvements in AUROC and Jaccard similarity of the GRN edge sets on 2 human and 5 mice datasets (Fig. 2). For all methods except CellOracle (CO), GRNITE provides a positive change in the AUROC. For the mDC and mESC datasets, use of GRNITE with CellOracle inferred target graph leads to a minor (*<* 0.01) decrease in the AUROC.

**Fig. 2:**
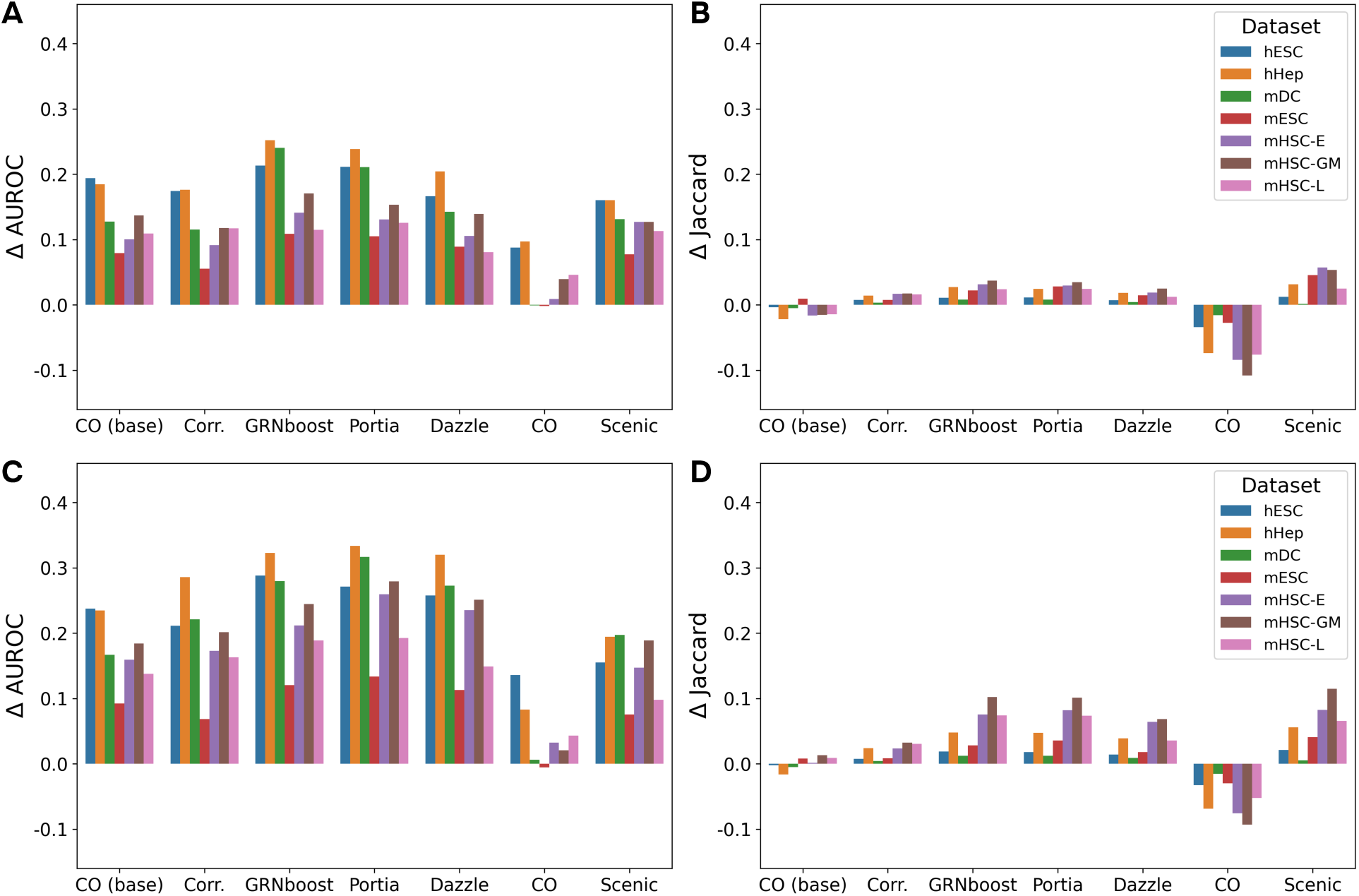
Change in performance from baseline methods to GRNITE. Change in the area under the receiver operating characteristic curve (*Δ*AUROC: A, C) and change in the Jaccard similarity of edge sets (*Δ*Jaccard: B, D) after our method has been applied to the target graph. Panels A and B correspond to BEELINE TF+500, and panels C and D correspond to BEELINE TF+1000 datasets.

Similarly, for all methods except CellOracle, GRNITE provides a moderate positive improvement in the Jaccard similarity of the inferred and true edge sets (Fig. 2). We note that the improvement in both AUROC and Jaccard similarity is higher in magnitude for BEELINE TF+1000 dataset (Fig. 2). For both BEELINE TF+500 and TF+1000 datasets all methods except CellOracle experience a positive change in F1-score, driven primarily by a large increase in recall (Supp. Fig. 8, 9). Notably, more conservative methods (CellOracle, SCENIC) which predict fewer edges, but have higher precision, lose some of their precision after application of GRNITE (Supp. Fig. 8, 9). Finally, ablations with respect to using a purely biological prior (**A**_bio_, Supp. Fig. 10, 11, 14, 15) or purely text embedding-based prior (**A**_emb_, Supp. Fig. 12, 13, 16, 17) show better performance from the combined prior across the majority of metrics.

#### GRouNdGAN datasets

For the GRouNdGAN simulated data, we observe a pattern of performance changes similar to the one on BEELINE. Specifically, all methods receive an improvement in AUROC, and CellOracle is the only method with consistently decreased Jaccard similarity for edge sets (Fig. 3).

**Fig. 3:**
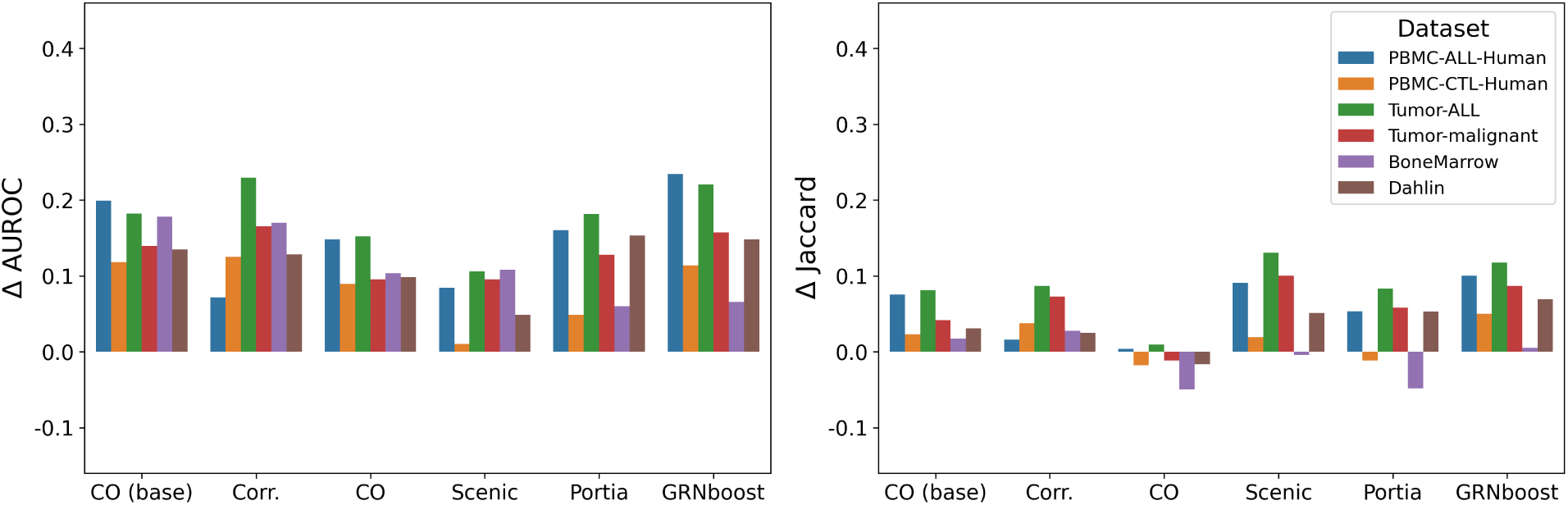
Change in performance from baseline methods to GRNITE on GRouNdGAN data. Change in the area under the receiver operating characteristic curve (*Δ*AUROC, left) and change in the Jaccard similarity of edge sets (*Δ*Jaccard) after our method has been applied to the target graph.

Notably, Portia also shows reduced Jaccard edge set similarity on the PBMC-CTL and Bone Marrow datasets. This matches the observation that these are the datasets on which GRNITE + Portia also shows a small reduction in precision (Supp. Fig. 18). Similarly to BEELINE, we observe a universal improvement in recall, and an improvement in F1-score for all the methods except CellOracle (Supp. Fig. 18). However, on these data, SCENIC experiences a large drop in precision after application of GRNITE, driven by a very low number of edge predictions in the original SCENIC method (*<*300 for all datasets, Supp. Fig. 18). Finally, analogous to the experiments on BEELINE, ablations with respect to using a purely biological prior (**A**_bio_, Supp. Fig. 19, 20) or purely text embedding-based prior (**A**_emb_, Supp. Fig. 21, 22) show better performance from the combined prior across the majority of metrics.

Furthermore, we produced five random subsamples, each containing 2,000 cells from the GRouNdGAN dataset, to assess the variance in the performance of GRNITE on subsets of the data. We note that the variance in the performance for all methods is small, with the exception of SCENIC, for which we observe larger variance between the AUROC on different cell subsets (Fig. 4). We omitted CellOracle base-graph and correlation network for these analyses, as these are not commonly employed inference methods, and are mainly relevant as baselines.

**Fig. 4:**
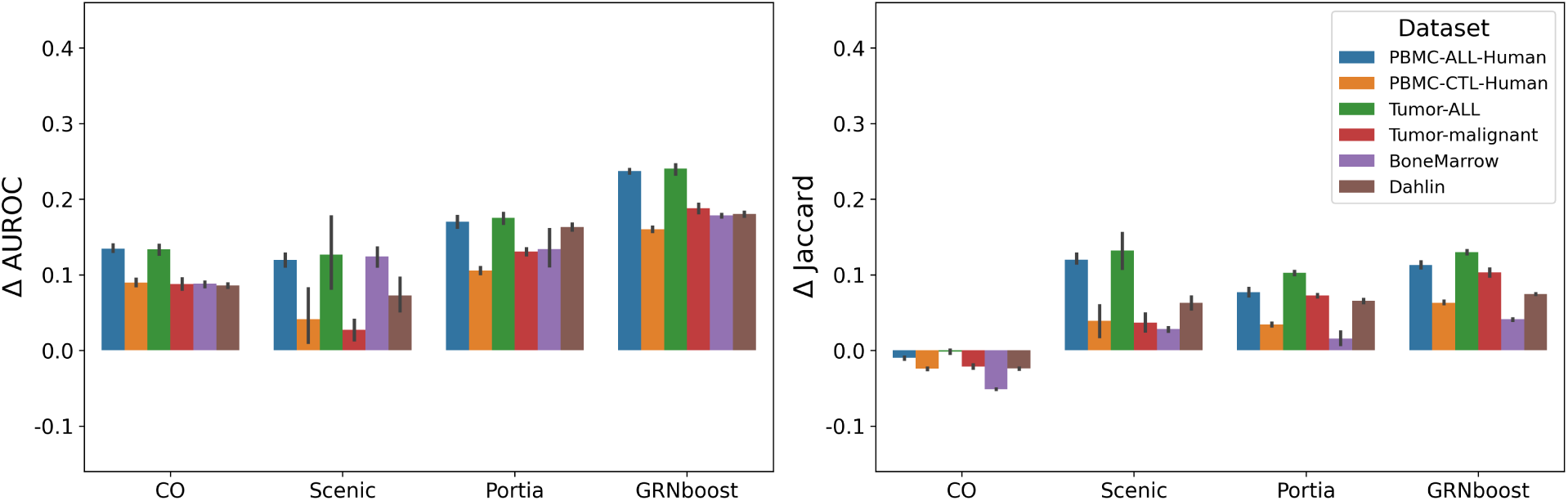
Change in performance from baseline methods to GRNITE on the subsamples of GRouNdGAN data. Change in the area under the receiver operating characteristic curve (*Δ*AUROC, left) and change in the Jaccard similarity of edge sets (*Δ*Jaccard) after our method has been applied to the target graph. Error bars indicate variance between 5 subsample runs.

### 3.3 Increasing inter-method consensus

Next, we evaluated the fraction of shared and unique edges predicted by each of the methods on the BEELINE and GRouNdGAN datasets. We measured the Jaccard similarity between all pairs of baseline methods, as well as the pairs of final GRNs that have been processed with GRNITE. We observe a large increase in pairwise Jaccard similarity of the edge sets after applying GRNITE (Fig. 5) for all datasets and methods evaluated.

**Fig. 5:**
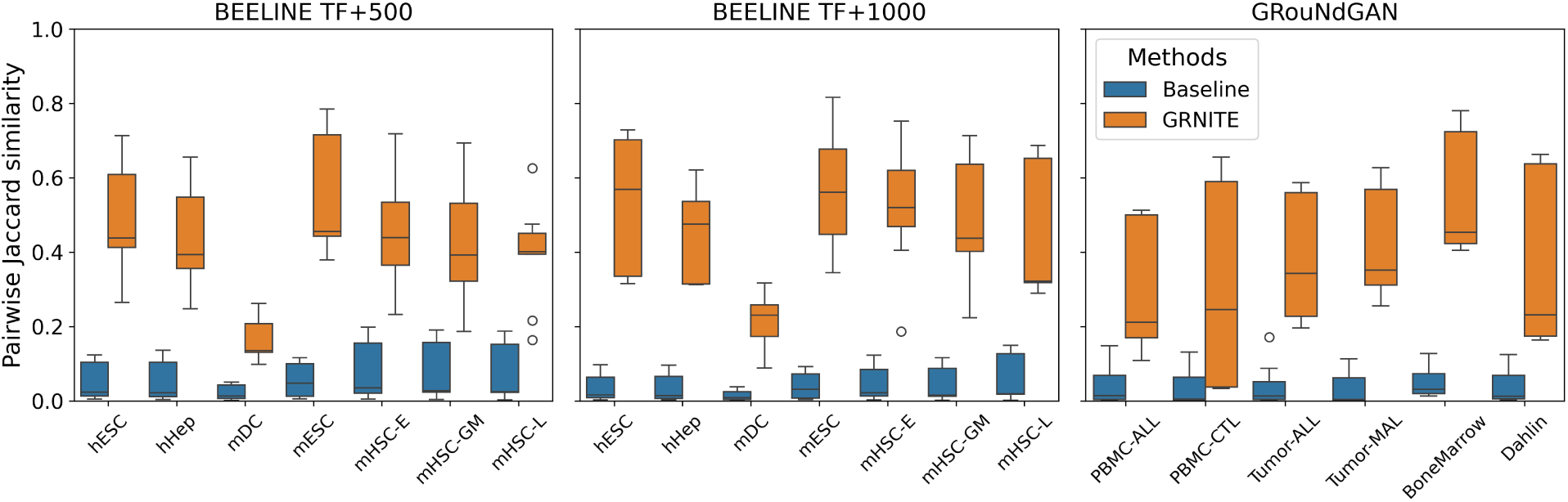
Pairwise Jaccard similarities between edge sets of GRNs inferred by the baseline methods (blue) and baseline + GRNITE (orange). Box plots show variation between all pairs of methods considered.

Furthermore, we note that the resulting edge set intersections between the pairs of methods have better AUROCs with respect to the ground truth (Fig. 6).

**Fig. 6:**
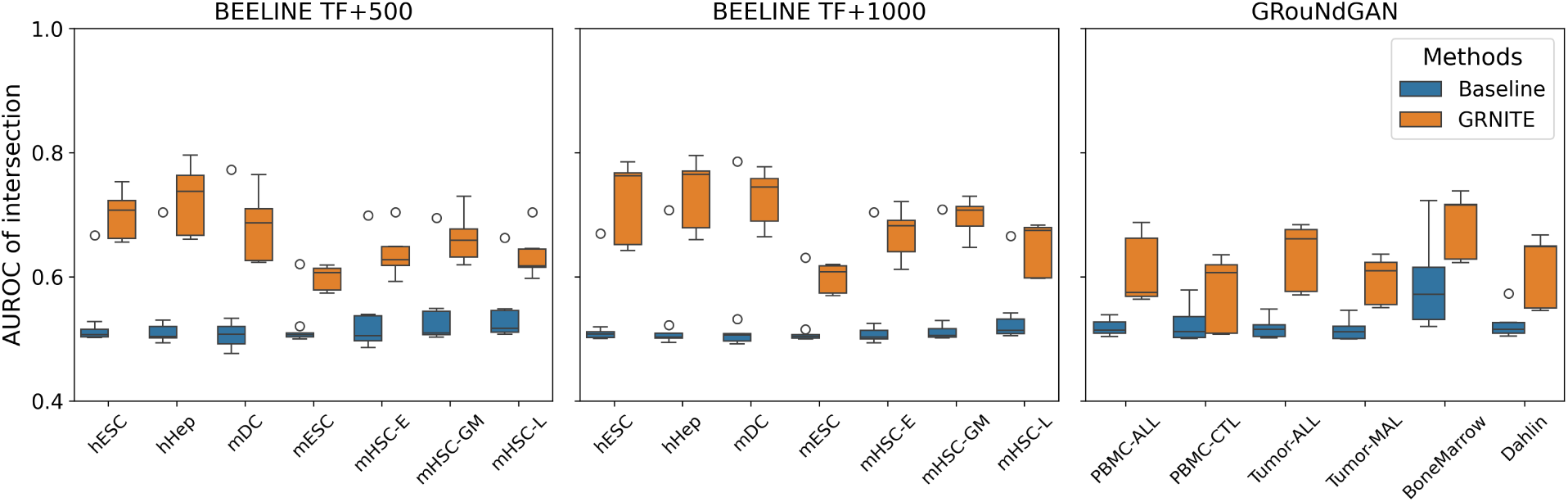
AUROC of the pairwise edge set intersections for GRNs inferred by the baseline methods (blue) and baseline + GRNITE (orange). Box plots show variation between all pairs of methods considered.

These observations indicate that applying GRNITE increases inter-method consensus, and the resulting consensus edge sets show an improved AUROC.

### 3.4 Evaluation of GRNITE on biological single-cell datasets

To evaluate GRNITE’s ability to enhance existing gene regulatory networks (GRNs) in recovering biologically meaningful regulatory interactions, we conducted a comparative evaluation across two biological single-cell datasets (HFD and PBMC, Section 2.4). For each dataset, GRNs were inferred using four baseline methods (GRNBoost, SCENIC, Portia, and CellOracle), followed by the generation of corresponding GRNITE-enhanced networks. We quantified performance using recall, defined as the fraction of ground-truth TF–target interactions recovered by each method (see Section 2.4 for details on curated ground-truth datasets). Across both the HFD and PBMC datasets, GRNITE consistently achieved substantial improvements in recall relative to the baselines (Fig. 7).

**Fig. 7:**
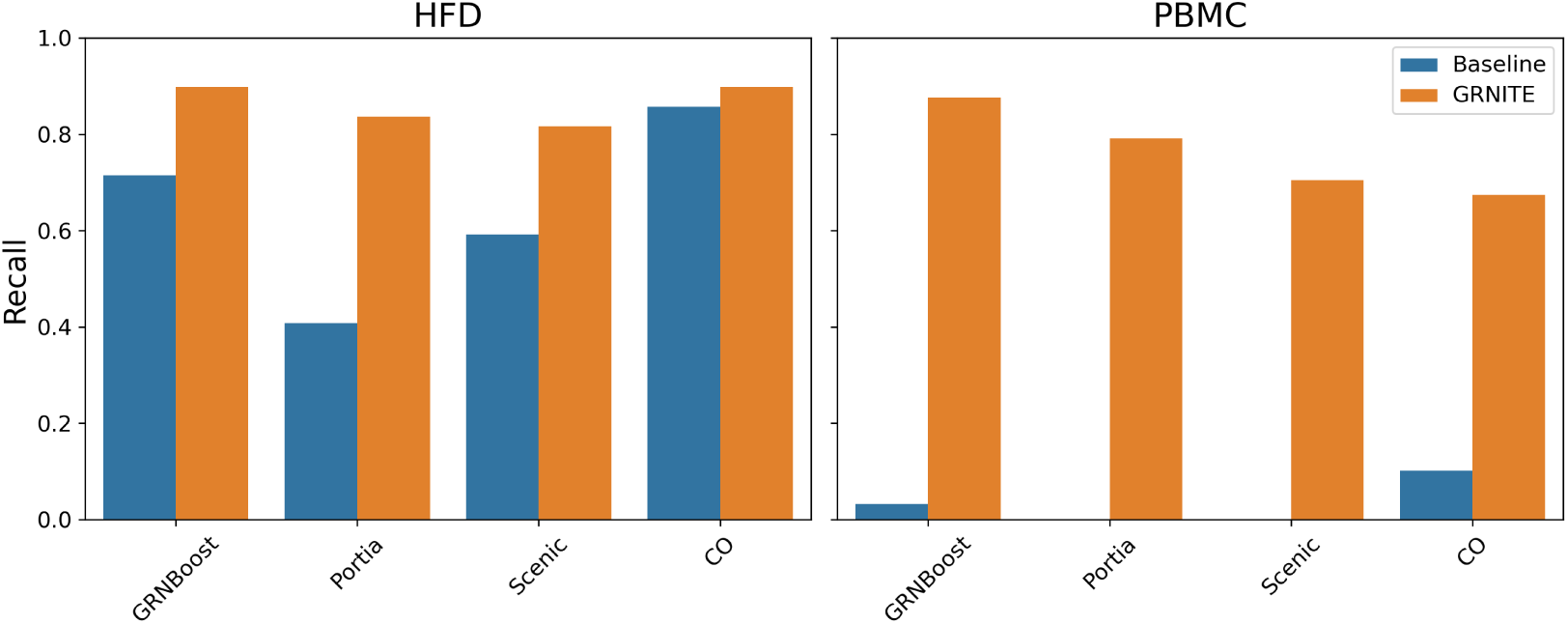
Change in recall of literature and experiment supported TF-target edges for baseline methods across the HFD (left) and PBMC (right) biological single-cell datasets.

To further characterize method-specific performance, we inferred GRNs for all baseline and GRNITE-enhanced models on the full 10x Genomics PBMC dataset and evaluated changes in AUROC and Jaccard similarity (Fig. 23). Notably, Portia and SCENIC exhibited reduced performance (lower precision/recall and recall) when evaluated against the PBMC GRouNdGAN ground truth (Fig. 24). These observations are consistent with the lower recall values observed for both methods in Fig. 7.

### 3.5 Computational Efficiency

We evaluate the total runtime required to infer GRNs using the original base methods and compare it with the time required to run GRNITE. GRNITE consists of three steps. The first two steps construct the text-and biology-informed prior graph (Section 2.1 and 2.2), which is a one-time preprocessing step and only needs to be computed once per dataset. In the third step (Section 2.3), this prior is combined with the output of any GRN inference method to refine and enhance its predictions. Since the prior is reusable, Step 3 can be applied efficiently to different GRN inference methods without recomputing Steps 1 and 2. Runtime results in seconds are reported in Table 2 for the BEELINE TF500 and TF1000 datasets. The reported times for Step 3 are averaged over all base methods.

**Table 2:**
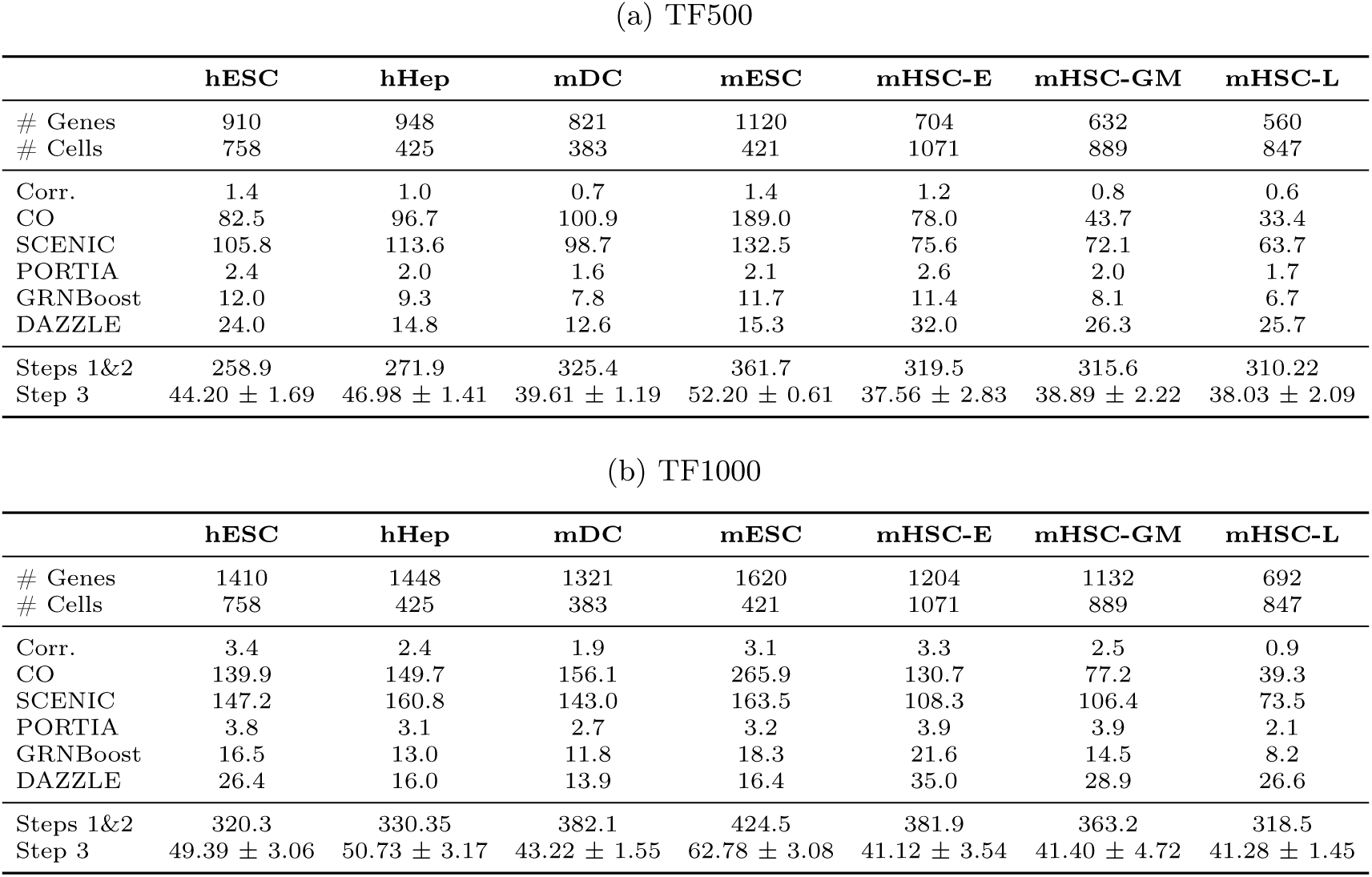
Runtime (in seconds) on BEELINE datasets for TF500 and TF1000. Each dataset reports number of genes, number of cells, baseline runtime of each method, Steps 1&2 prior construction time, and GRNITE refinement time in Step 3.

| (a) TF500 |  |  |  |  |  |  |  |
| --- | --- | --- | --- | --- | --- | --- | --- |
|  | hESC | hHep | mDC | mESC | mHSC-E | mHSC-GM | mHSC-L |
| # Genes | 910 | 948 | 821 | 1120 | 704 | 632 | 560 |
| # Cells | 758 | 425 | 383 | 421 | 1071 | 889 | 847 |
| Corr. | 1.4 | 1.0 | 0.7 | 1.4 | 1.2 | 0.8 | 0.6 |
| CO | 82.5 | 96.7 | 100.9 | 189.0 | 78.0 | 43.7 | 33.4 |
| SCENIC | 105.8 | 113.6 | 98.7 | 132.5 | 75.6 | 72.1 | 63.7 |
| PORTIA | 2.4 | 2.0 | 1.6 | 2.1 | 2.6 | 2.0 | 1.7 |
| GRNBoost | 12.0 | 9.3 | 7.8 | 11.7 | 11.4 | 8.1 | 6.7 |
| DAZZLE | 24.0 | 14.8 | 12.6 | 15.3 | 32.0 | 26.3 | 25.7 |
| Steps 1&2 | 258.9 | 271.9 | 325.4 | 361.7 | 319.5 | 315.6 | 310.22 |
| Step 3 | 44.20 ± 1.69 | 46.98 ± 1.41 | 39.61 ± 1.19 | 52.20 ± 0.61 | 37.56 ± 2.83 | 38.89 ± 2.22 | 38.03 ± 2.09 |

| (b) TF1000 |  |  |  |  |  |  |  |
| --- | --- | --- | --- | --- | --- | --- | --- |
|  | hESC | hHep | mDC | mESC | mHSC-E | mHSC-GM | mHSC-L |
| # Genes | 1410 | 1448 | 1321 | 1620 | 1204 | 1132 | 692 |
| # Cells | 758 | 425 | 383 | 421 | 1071 | 889 | 847 |
| Corr. | 3.4 | 2.4 | 1.9 | 3.1 | 3.3 | 2.5 | 0.9 |
| CO | 139.9 | 149.7 | 156.1 | 265.9 | 130.7 | 77.2 | 39.3 |
| SCENIC | 147.2 | 160.8 | 143.0 | 163.5 | 108.3 | 106.4 | 73.5 |
| PORTIA | 3.8 | 3.1 | 2.7 | 3.2 | 3.9 | 3.9 | 2.1 |
| GRNBoost | 16.5 | 13.0 | 11.8 | 18.3 | 21.6 | 14.5 | 8.2 |
| DAZZLE | 26.4 | 16.0 | 13.9 | 16.4 | 35.0 | 28.9 | 26.6 |
| Steps 1&2 | 320.3 | 330.35 | 382.1 | 424.5 | 381.9 | 363.2 | 318.5 |
| Step 3 | 49.39 ± 3.06 | 50.73 ± 3.17 | 43.22 ± 1.55 | 62.78 ± 3.08 | 41.12 ± 3.54 | 41.40 ± 4.72 | 41.28 ± 1.45 |

## 4 Discussion

In this study, we introduced GRNITE, a flexible meta-method designed to enhance the performance of any scRNA-seq–based GRN inference algorithm, with a particular emphasis on improving recall. Using both BEELINE and GRouNdGAN benchmarking datasets, we demonstrated that GRNITE substantially increases the recall of existing GRN predictions. Furthermore, for some methods GRNITE also improves precision, while for more conservative methods it slightly reduces precision while greatly expanding the number of correctly identified TF-terget interactions. Additionally, we showed that GRNITE strengthens inter-method consensus and yields higher AUROC values when evaluated on consensus edge sets, highlighting its ability to reconcile and refine heterogeneous GRN outputs. This feature is particularly appealing for biological data, where analyses performed by different methods often yield vastly different inference results.

Our analyses on biological single-cell datasets illustrate that GRNITE not only boosts recall and inter-method consensus, but also enhances the ability of baseline methods to recover biologically meaningful TF–target interactions. We also indicate GRNITE’s ability to improve overall performance, characterized by improved AUROC and Jaccard similarity, of existing GRN inference methods over the biological PBMC dataset.

We observe that the time required to run GRNITE (particularly Step 3) is comparable to the cost of running the baseline methods evaluated in this study. This implies that processing of the inferred GRN with GRNITE does not inquire a signficant computational cost, which makes it an attractive extra step in GRN inference and analysis pipelines. In comparison, Steps 1 and 2 are comparatively more time-consuming than Step 3, because it involves constructing the prior graph by heuristic matching of biological interactions to the gene set of interest. However, these steps can be performed in parallel with the baseline GRN inference.

Collectively, our findings position GRNITE as a robust and generalizable enhancement layer for state-of-the-art scRNA-seq GRN inference tools. By improving recall with only modest precision trade-offs, GRNITE enables existing methods to more reliably capture the underlying regulatory architecture of new datasets, ultimately advancing efforts to reconstruct accurate GRNs from single-cell transcriptomic data.

## Acknowledgments

This work was in part supported by the NSF grants DMS/NIGMS-2153704, DBI-2030604, IIS-2106837, EF-2126387, and CCF-2340481, and Rice’s Ken Kennedy Institute.

## A Supplementary figures and tables

### A.1 Precision, Recall and F1 changes from baseline methods on BEELINE

**Fig. 8:**
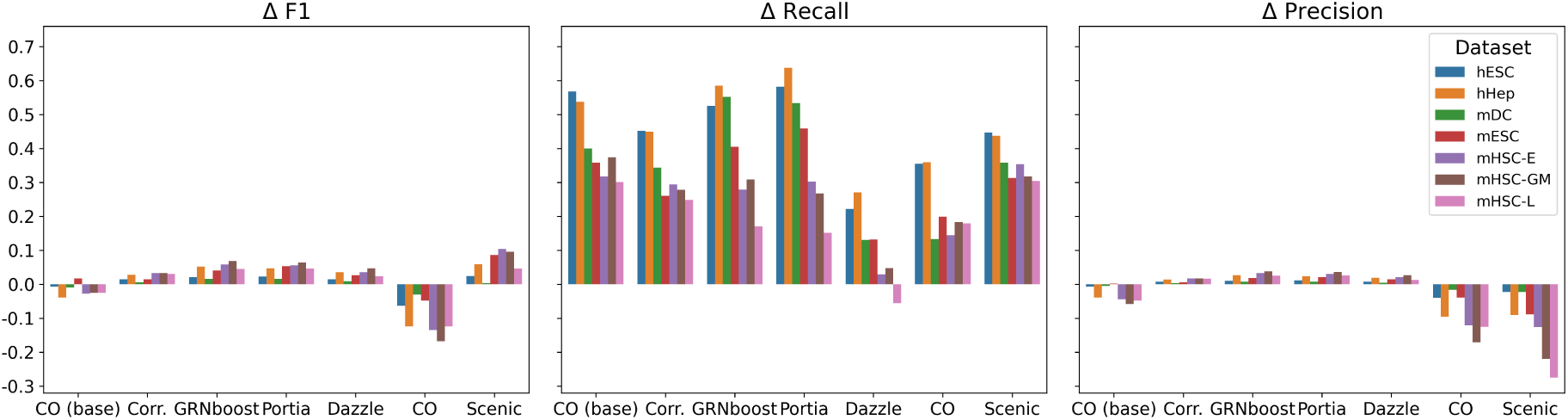
Change in the F1 (*Δ*F1, left), Recall (*Δ*Recall, middle) and Precision (*Δ*Precision) between the baseline GRNs and GRNITE enhanced GRNs across BEELINE TF500 datasets.

**Fig. 9:**
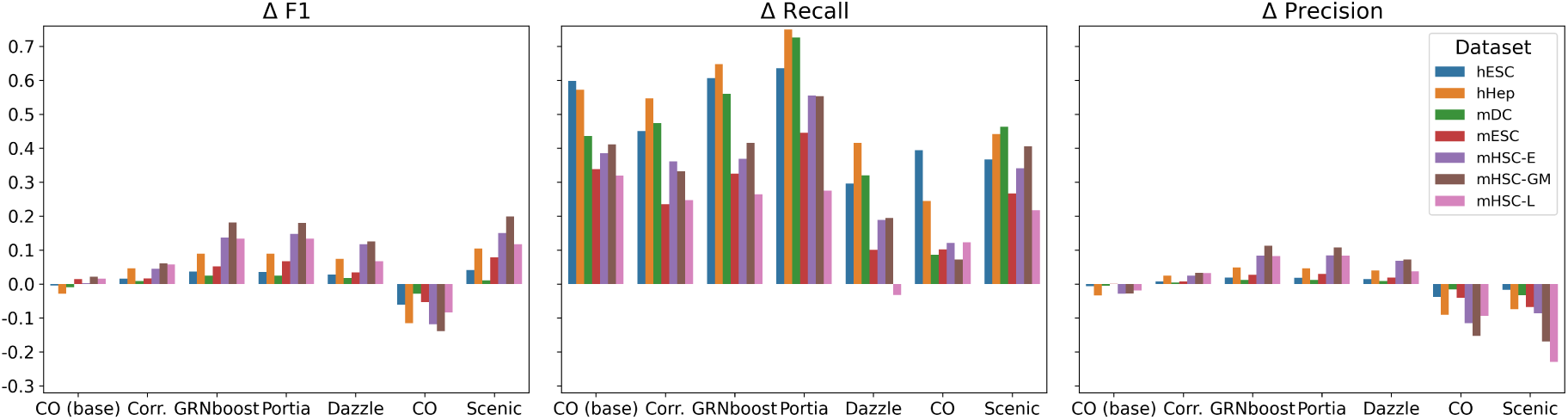
Change in the F1 (*Δ*F1, left), Recall (*Δ*Recall, middle) and Precision (*Δ*Precision) between the baseline GRNs and GRNITE enhanced GRNs across BEELINE TF1000 datasets.

### A.2 Prior graph ablation results for BEELINE TF+500

**Fig. 10:**
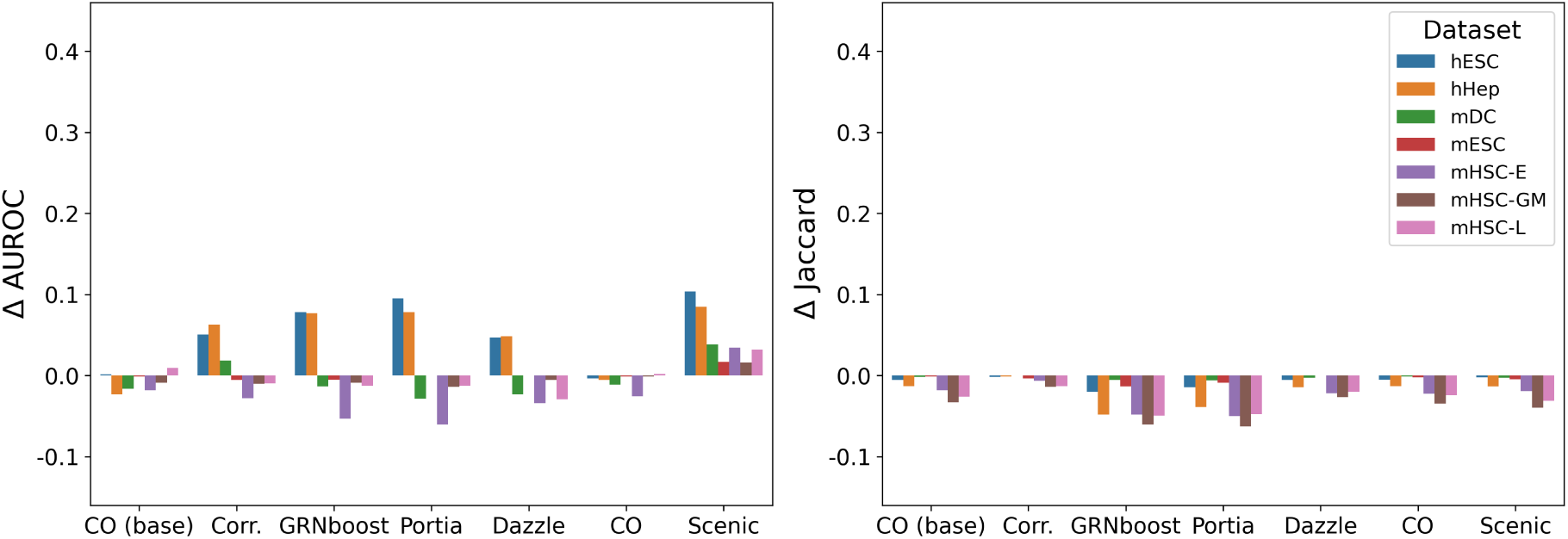
Change in the area under the receiver operating characteristic curve (*Δ*AUROC, left) and change in the Jaccard similarity (*Δ*Jaccard) on using the prior graph *A*_bio_ for BEELINE TF500 datasets.

**Fig. 11:**
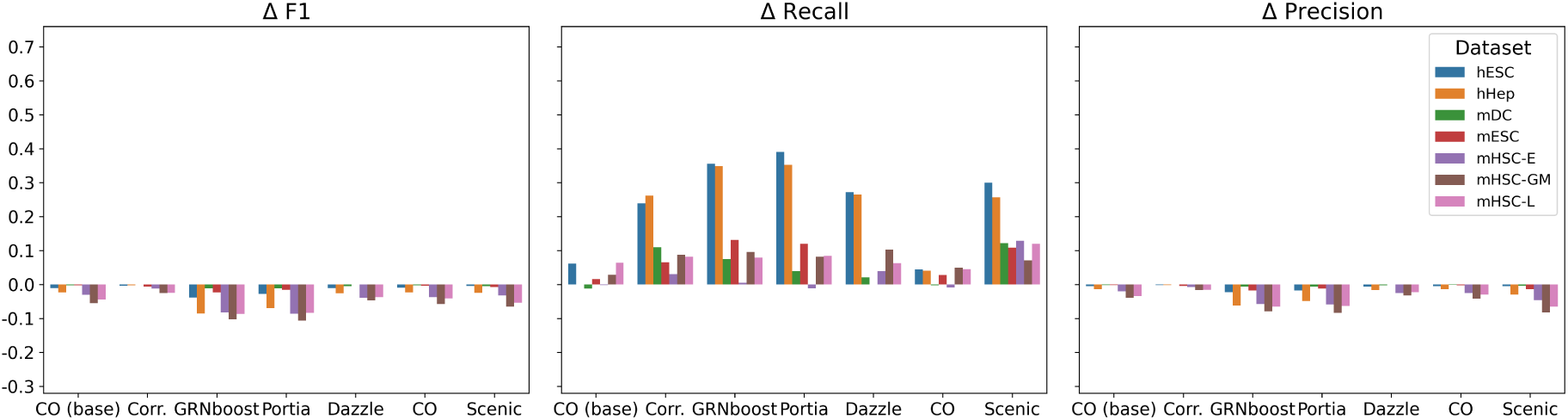
Change in the the F1 (*Δ*F1, left), Recall (*Δ*Recall, middle) and Precision (*Δ*Precision) on using the prior graph *A*_bio_ for BEELINE TF500 datasets.

**Fig. 12:**
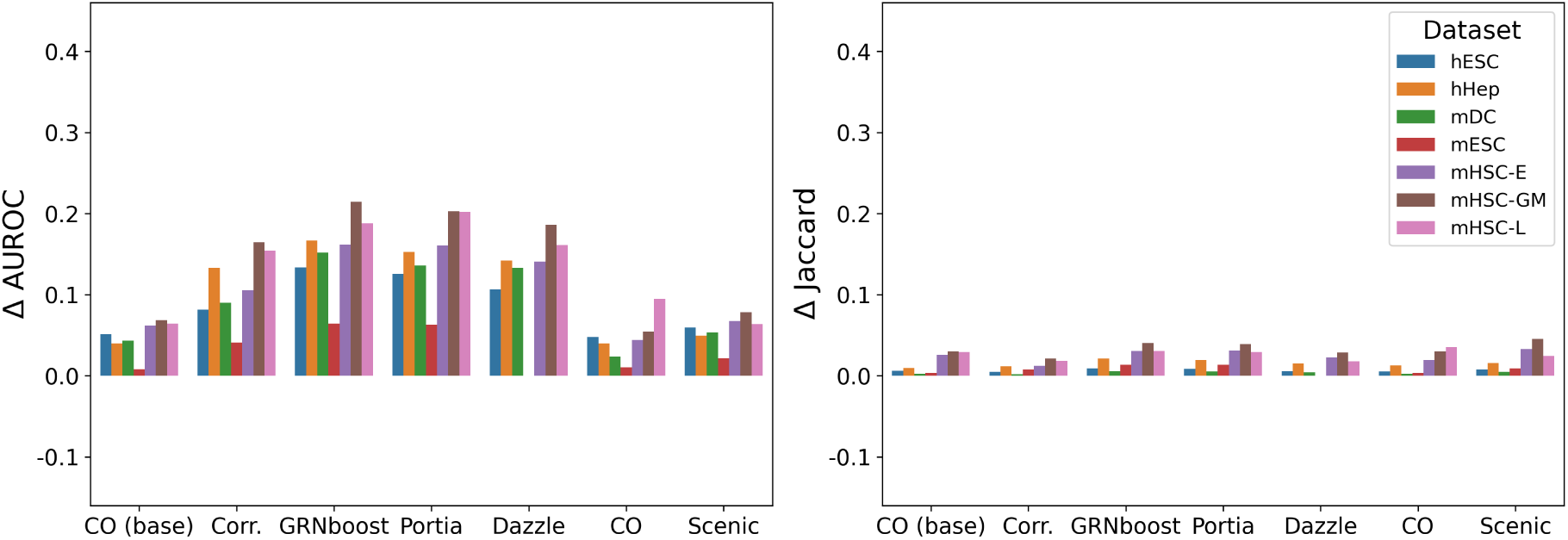
Change in the area under the receiver operating characteristic curve (*Δ*AUROC, left) and change in the Jaccard similarity (*Δ*Jaccard) on using the prior graph *A*_emb_ for BEELINE TF500 datasets.

**Fig. 13:**
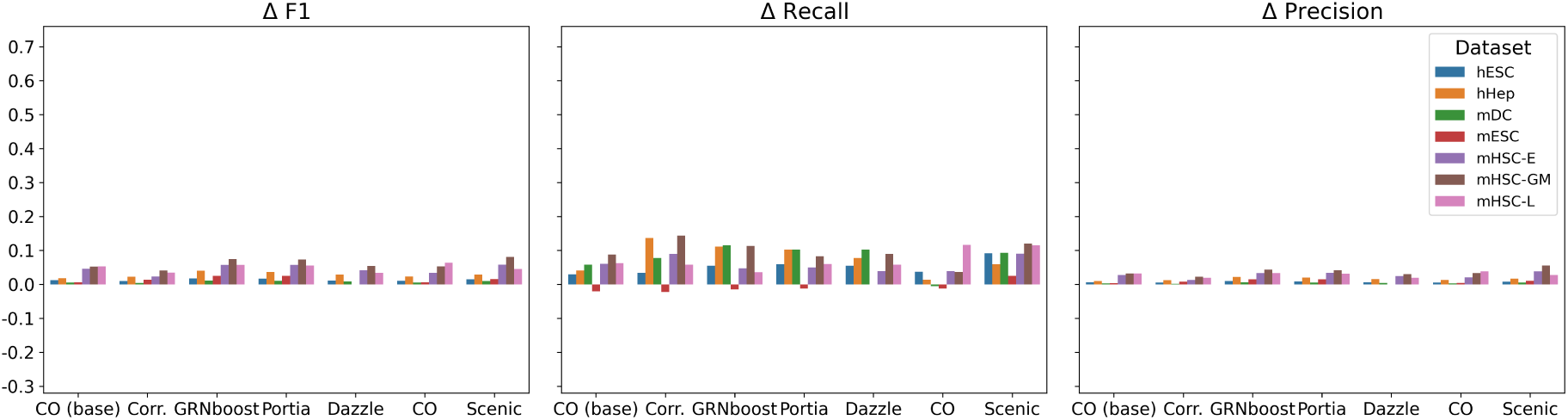
Change in the the F1 (*Δ*F1, left), Recall (*Δ*Recall, middle) and Precision (*Δ*Precision) on using the prior graph *A*_emb_ for BEELINE TF500 datasets.

### A.3 Prior graph ablation results for BEELINE TF+1000

**Fig. 14:**
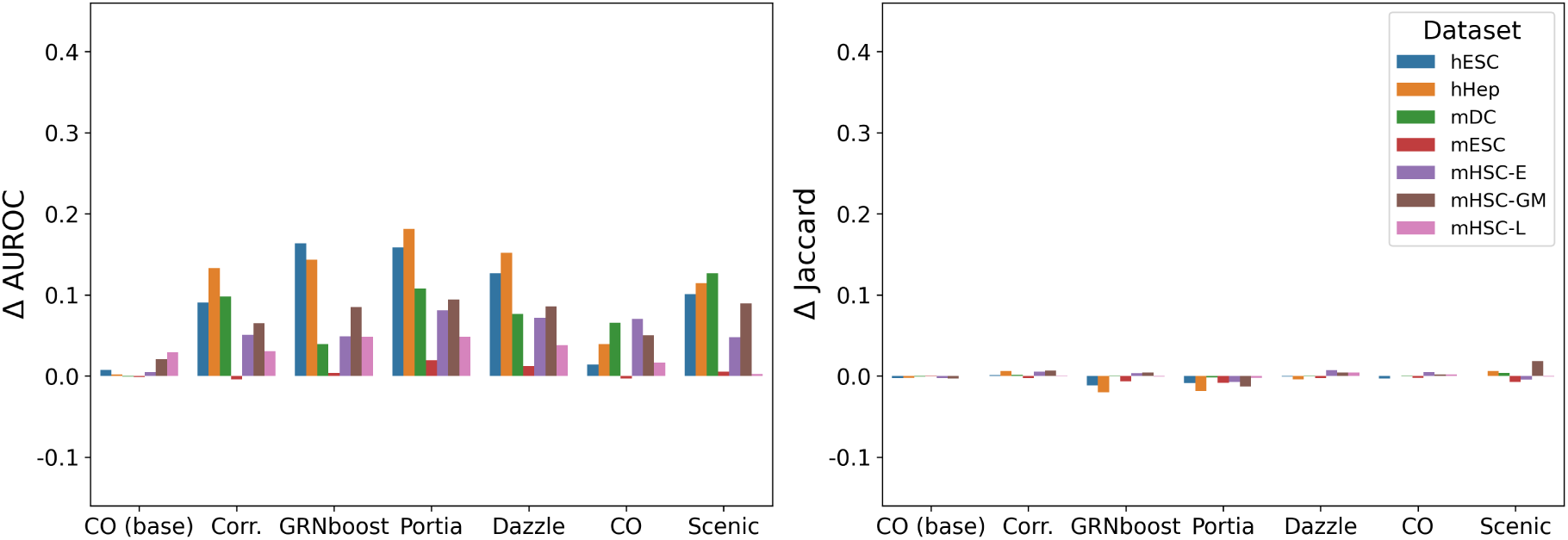
Change in the area under the receiver operating characteristic curve (*Δ*AUROC, left) and change in the Jaccard similarity (*Δ*Jaccard) on using the prior graph *A*_bio_ for BEELINE TF1000 datasets.

**Fig. 15:**
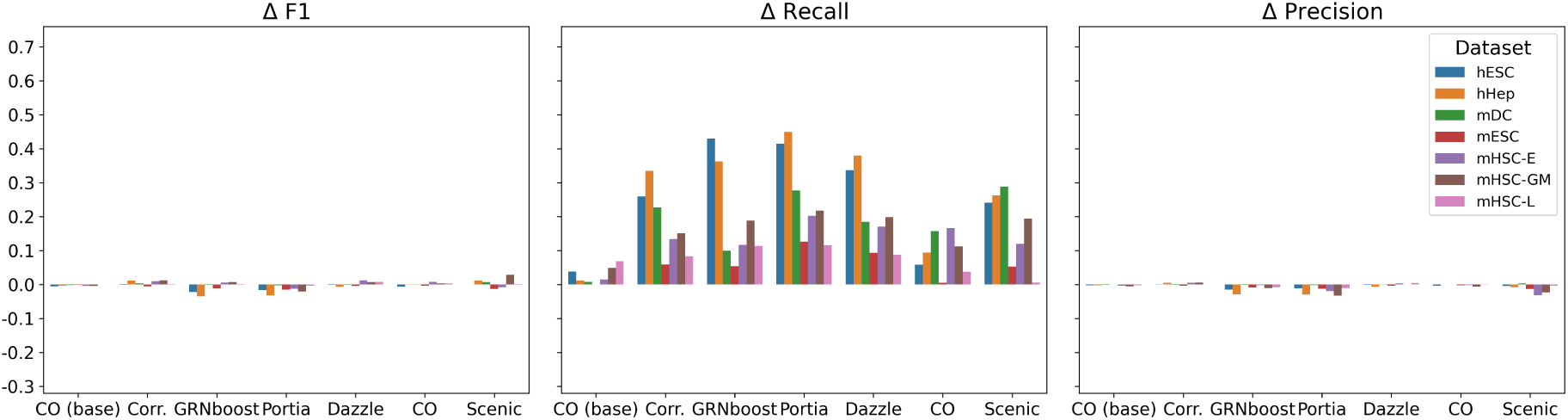
Change in the the F1 (*Δ*F1, left), Recall (*Δ*Recall, middle) and Precision (*Δ*Precision) on using the prior graph *A*_bio_ for BEELINE TF1000 datasets.

**Fig. 16:**
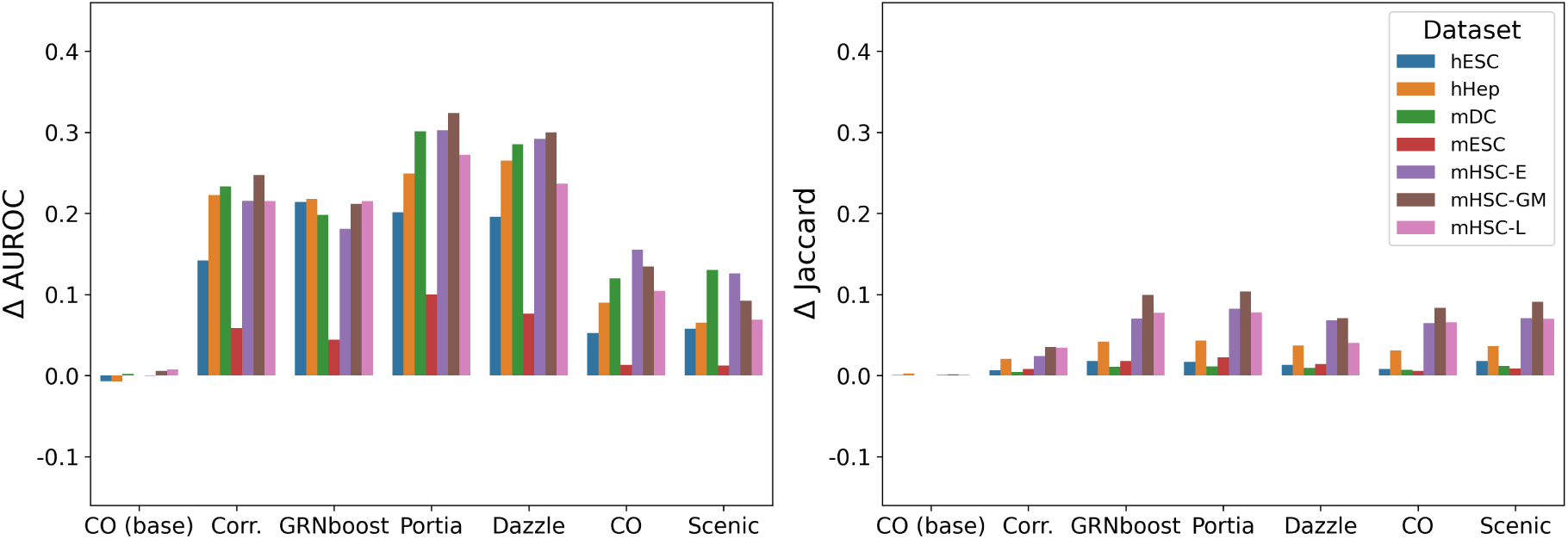
Change in the area under the receiver operating characteristic curve (*Δ*AUROC, left) and change in the Jaccard similarity (*Δ*Jaccard) on using the prior graph *A*_emb_ for BEELINE TF1000 datasets.

**Fig. 17:**
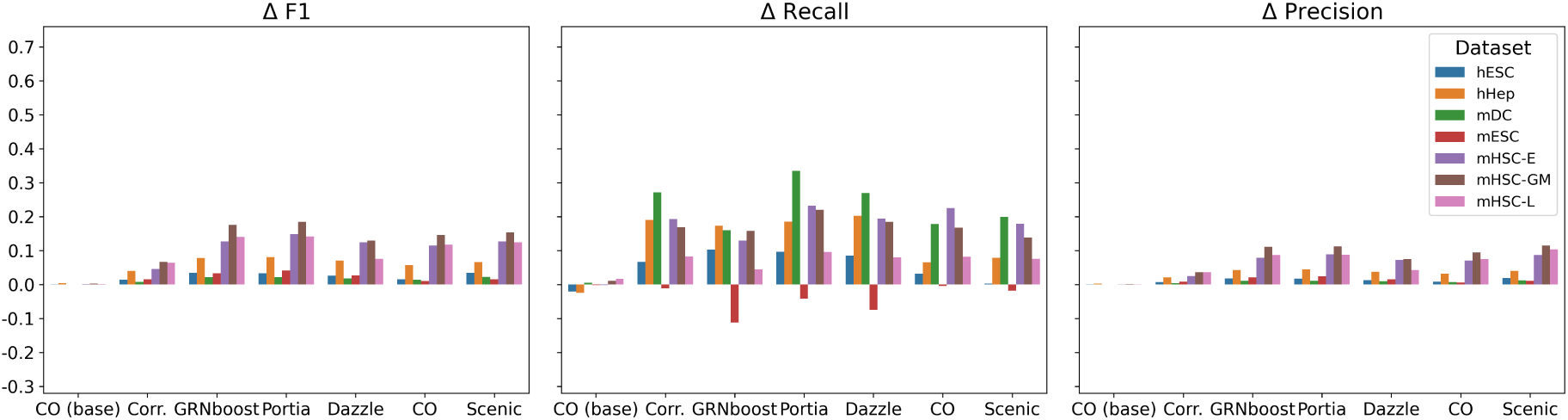
Change in the the F1 (*Δ*F1, left), Recall (*Δ*Recall, middle) and Precision (*Δ*Precision) on using the prior graph *A*_emb_ for BEELINE TF1000 datasets.

### A.4 Precision, Recall and F1 changes from baseline methods on GRouNdGAN

**Fig. 18:**
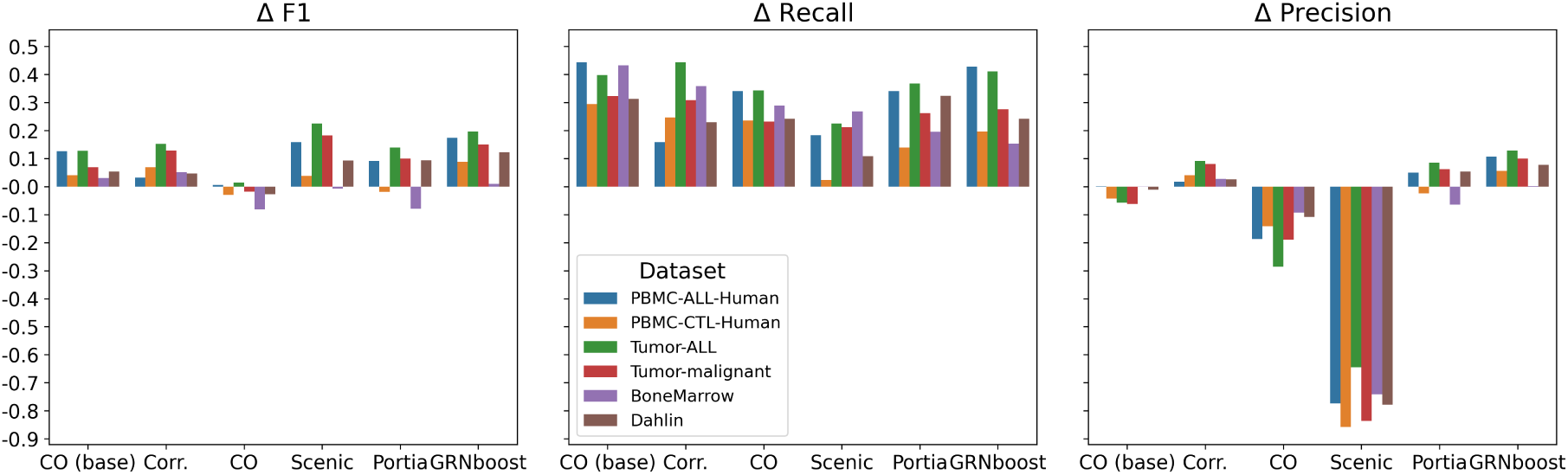
Change in the F1 (*Δ*F1, left), Recall (*Δ*Recall, middle) and Precision (*Δ*Precision) between the baseline GRNs and GRNITE enhanced GRNs across GRouNdGAN datasets.

### A.5 Prior graph ablation results for GRouNdGAN

**Fig. 19:**
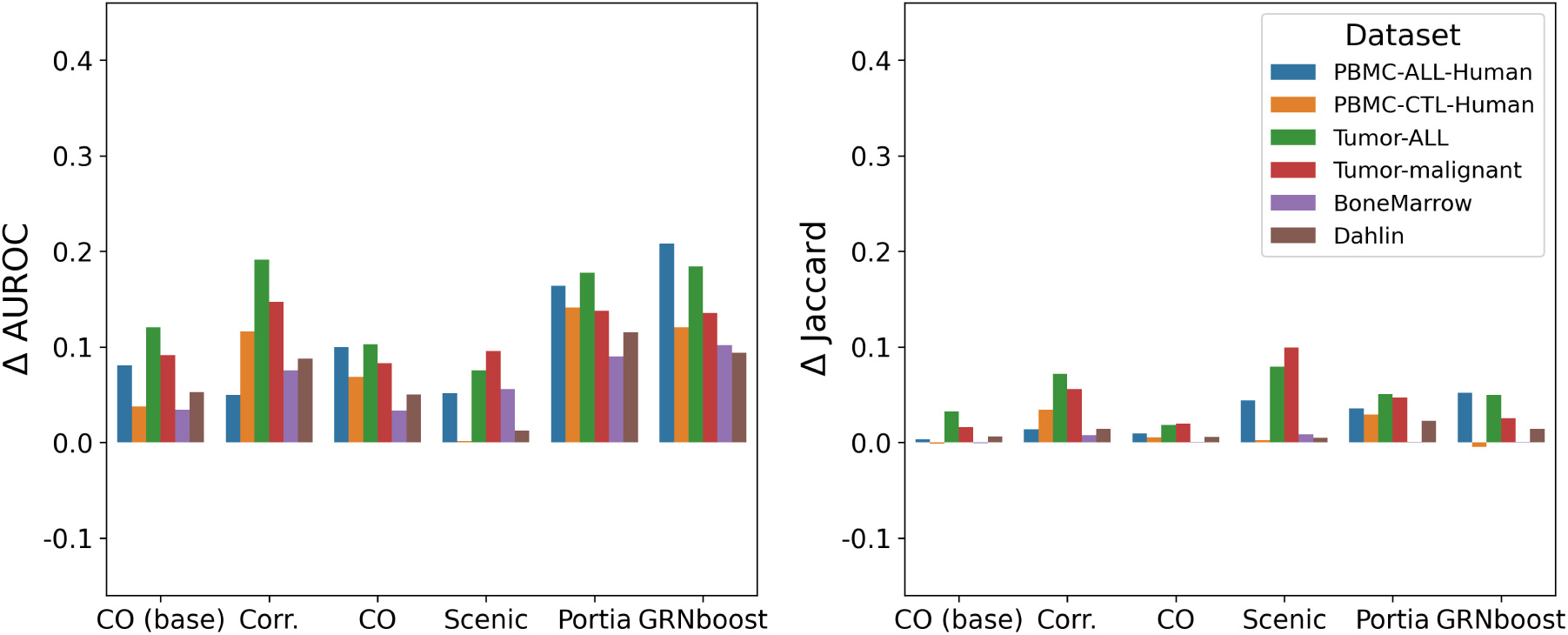
Change in the area under the receiver operating characteristic curve (*Δ*AUROC, left) and change in the Jaccard similarity (*Δ*Jaccard) on using the prior graph *A*_bio_ for GRouNdGAN datasets.

**Fig. 20:**
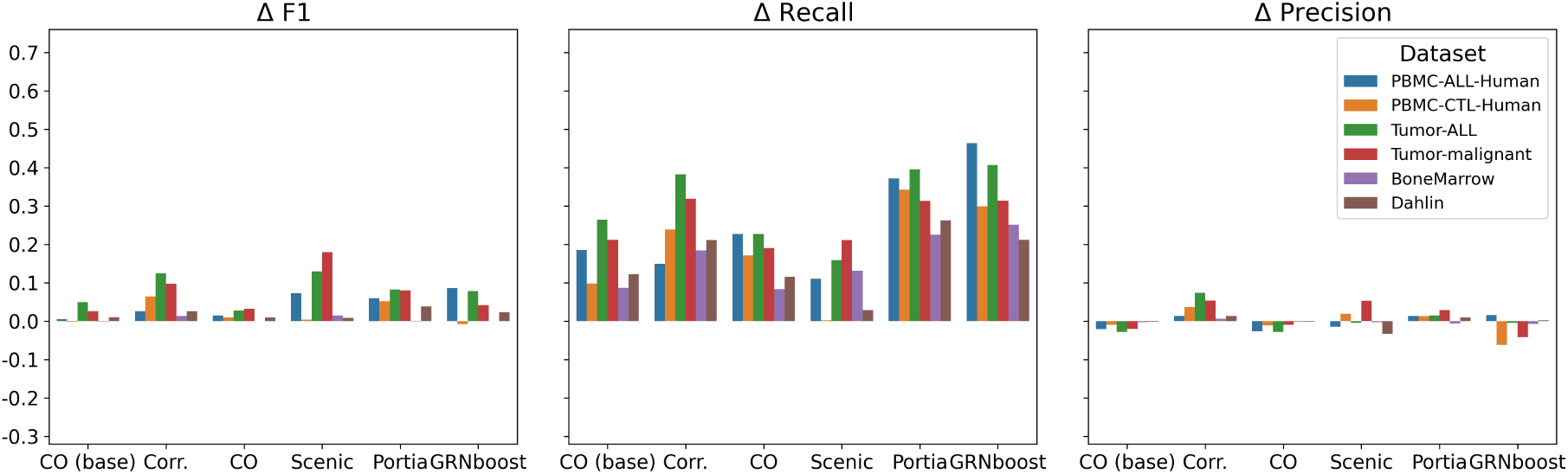
Change in the the F1 (*Δ*F1, left), Recall (*Δ*Recall, middle) and Precision (*Δ*Precision) on using the prior graph *A*_bio_ for GRouNdGAN datasets.

**Fig. 21:**
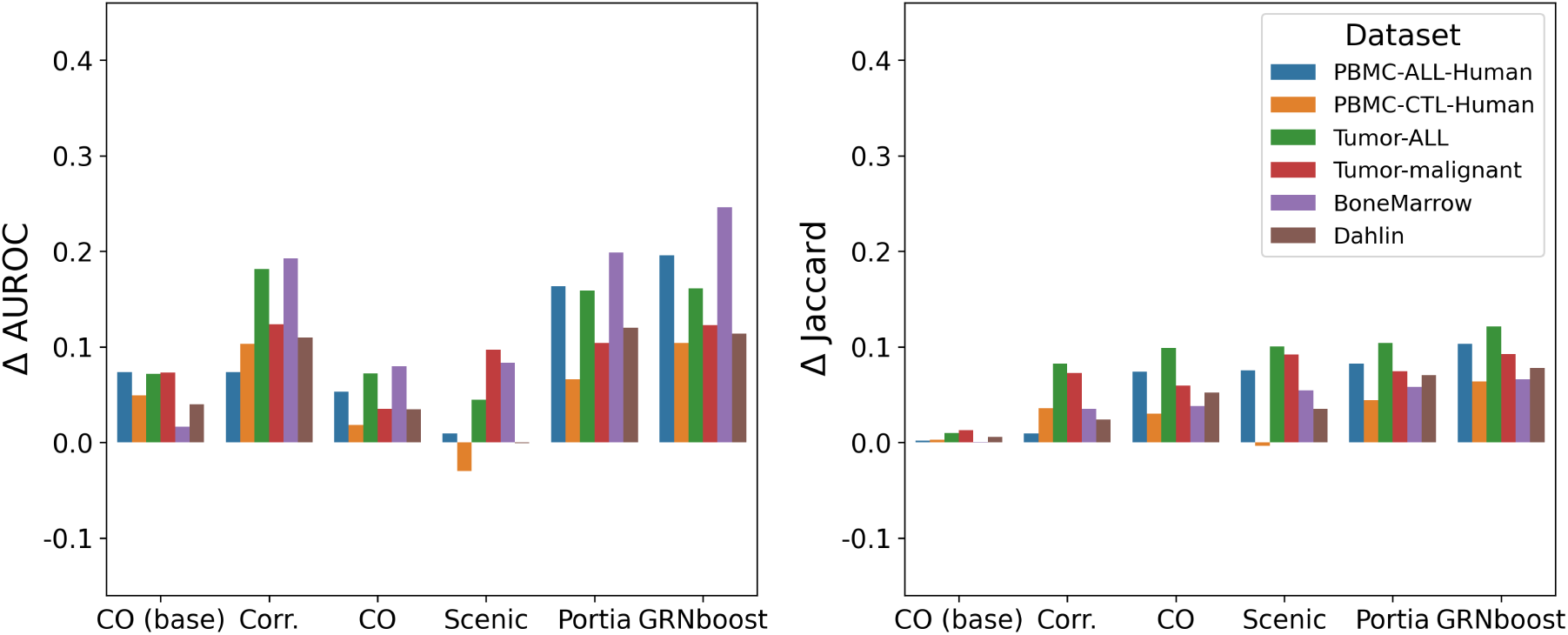
Change in the area under the receiver operating characteristic curve (*Δ*AUROC, left) and change in the Jaccard similarity (*Δ*Jaccard) on using the prior graph *A*_emb_ for GRouNdGAN datasets.

**Fig. 22:**
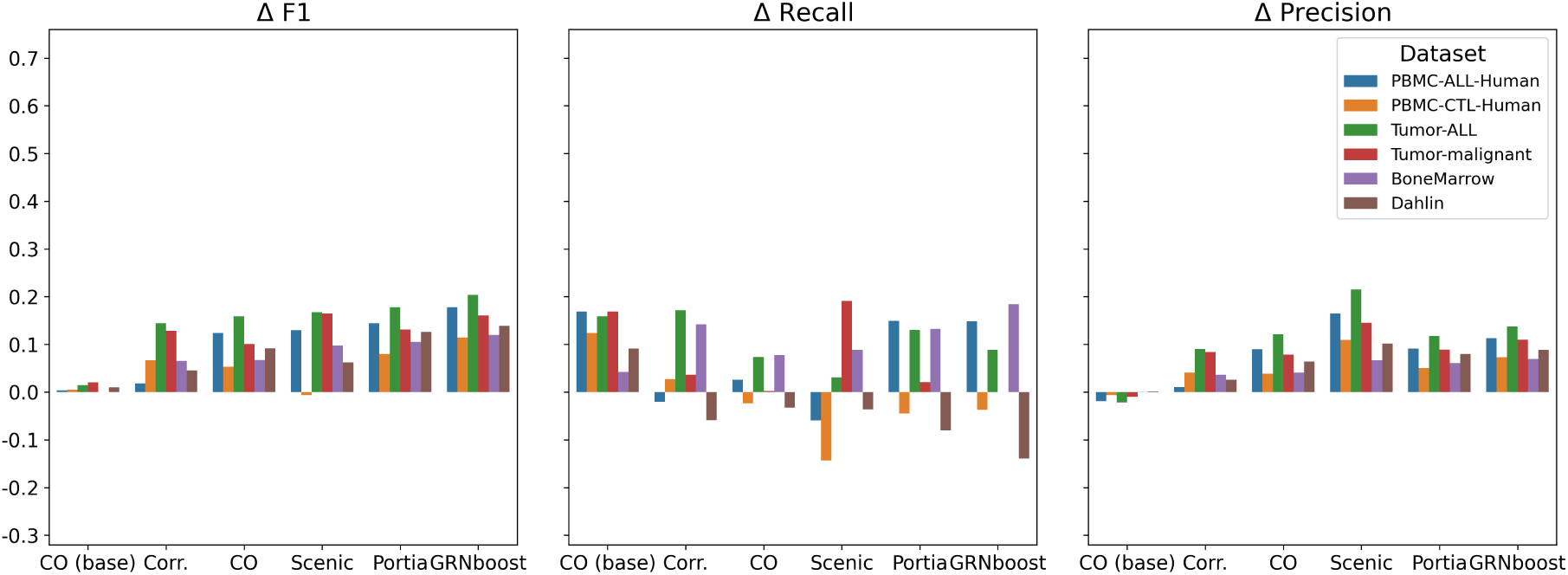
Change in the the F1 (*Δ*F1, left), Recall (*Δ*Recall, middle) and Precision (*Δ*Precision) on using the prior graph *A*_emb_ for GRouNdGAN datasets.

### A.6 AUROC, Jaccard, Precision and Recall changes from baseline methods on biological PBMC

**Fig. 23:**
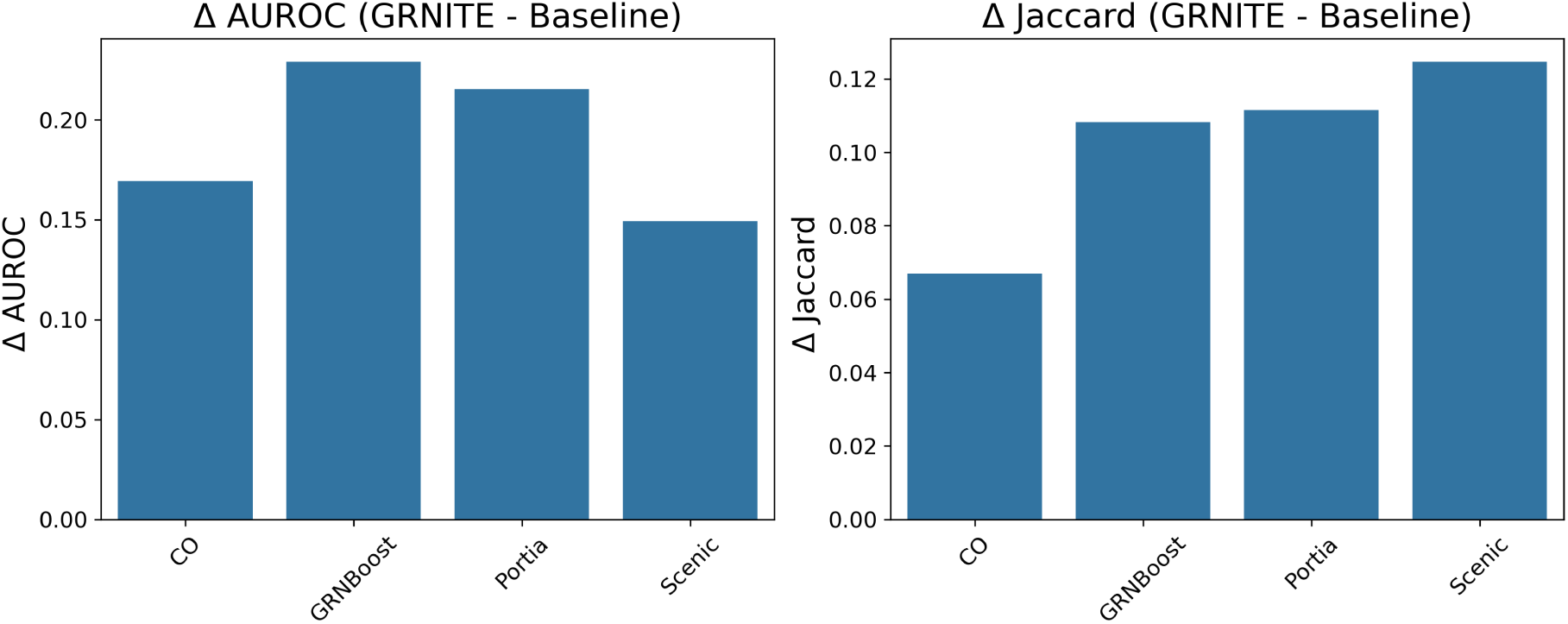
Change in the area under the receiver operating characteristic curve (*Δ*AUROC, left) and change in Jaccard similarity (*Δ*Jaccard) between the the baseline and enhanced GRNITE GRN.

**Fig. 24:**
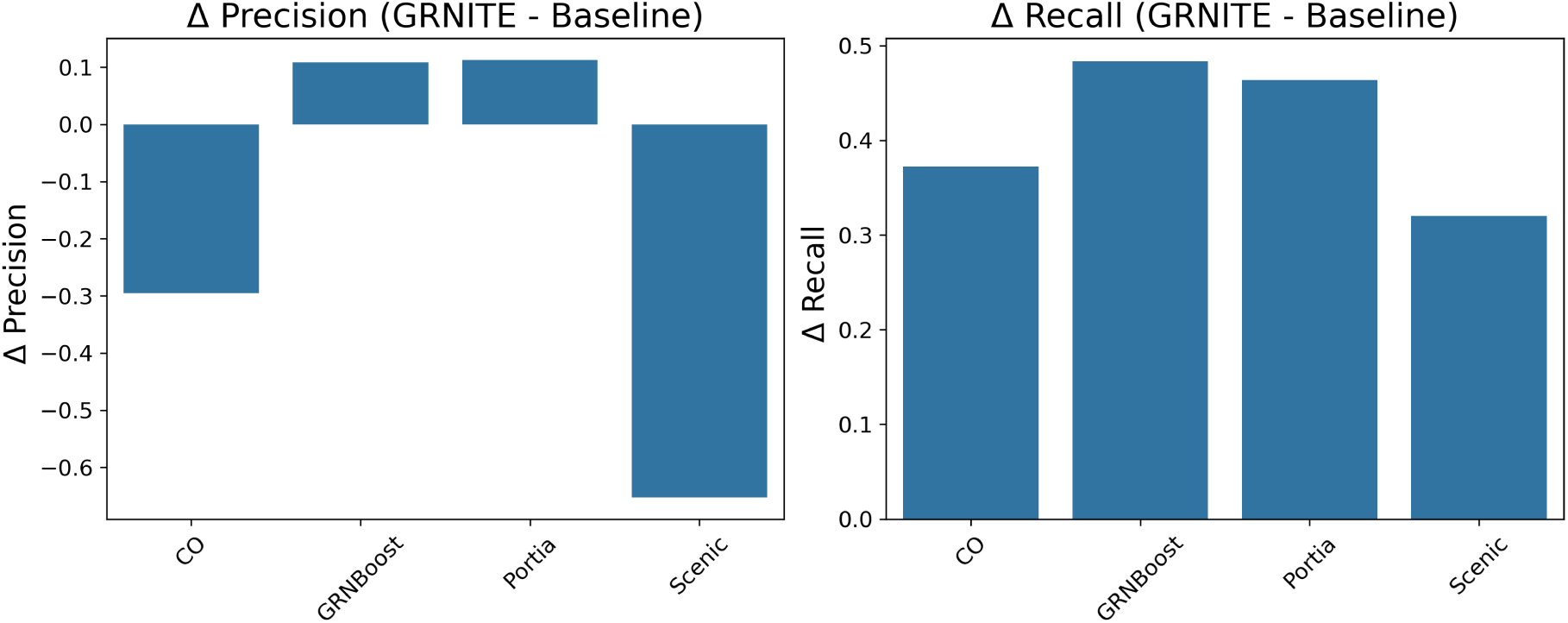
Change in Precision (*Δ*Precision, left) and change in Recall (*Δ*Recall) between the the baseline and enhanced GRNITE GRN.

## B Hyper-parameters

The hyperparameters used in GRNITE are kept consistent across all datasets and steps unless otherwise specified. We use the Adam optimizer [13] with a learning rate of *lr* = 0.01 and train the model for 5000 epochs. The graph autoencoder employs a GNN encoder with two hidden layers of dimensions [32, 32], each followed by a nonlinear relu activation. The feature dimensionality is reduced to *n*_low_ = 100 using TruncatedSVD prior to training for gene expressions in Step 1. During optimization, negative edge samples are generated with a ratio of neg_multiplier=1 relative to positive edges. The loss function in Step 2 is a weighted combination of the target and prior losses, with trade-off coefficient *β* = 0.5. Finally, *τ*_1_ is set to the mean of all values in **W**_emb_ in Section 2.1 and *τ*_2_ and *τ*_3_ are set to 0.5 in Sections 2.2 and 2.3.

## C Compute resources

All experiments were conducted on a server running Ubuntu 20.04.6 LTS, equipped with an AMD EPYC 7742 64-Core Processor and an NVIDIA A100-SXM4-80GB GPU with 80GB of memory. For model development, we utilized PyTorch version 2.5.1, along with PyTorch Geometric version 2.6.1, which also served as the source for all datasets used in our study.

